# A morphological comparison of the caudal rami of the superior temporal sulcus in humans, chimpanzees, and other great apes

**DOI:** 10.1101/2025.07.16.665137

**Authors:** Reyansh N. Sathishkumar, Ethan H. Willbrand, Priyanka Nanayakkara, Willa I. Voorhies, Yi-Heng Tsai, Thomas Gagnant, William D. Hopkins, Chet C. Sherwood, Kevin S. Weiner

## Abstract

For centuries, anatomists have charted the folding patterns of the sulci of the cerebral cortex in primates. Improvements in neuroimaging technologies over the past decades have led to advancements in understanding of the sulcal organization of the human cerebral cortex, yet comparisons to chimpanzees, one of humans’ closest extant phylogenetic relatives, remain to be performed in many regions, such as superior temporal cortex. For example, while several posterior branches, or rami, of the superior temporal sulcus (STS) have been identified in great apes since the late 1800s, no study has yet to comprehensively identify and quantitatively compare these rami across species. To fill this gap in knowledge, in the present study, we defined the three caudal branches of the STS (cSTS) in 72 human and 29 chimpanzee brains (202 total hemispheres) and then extracted and compared the morphological (depth and surface area) properties of these sulci. We report three main findings. First, modern methods replicate classic findings that three rami of the posterior STS are unique to the hominid lineage (i.e., humans and great apes). Second, normalizing for brain size, the cSTS rami were relatively deeper in chimpanzees compared to humans. Third, the cSTS branches were relatively larger in surface area in humans compared to chimpanzees. Finally, we share probabilistic predictions of the cSTS to guide the identification of these sulci in future studies. Altogether, these findings bridge the gap between historic qualitative observations and modern quantitative measurements in a part of the brain that has expanded substantially throughout evolution and that is involved in human-specific aspects of cognition.

## Introduction

Exploring the similarities and differences in brain structure among species is of major interest across neurobiological subdisciplines. For centuries, a central goal has been to determine which features of the central nervous system (CNS) are specific to humans. A feature of the CNS that offers unique utility to address this aim is the folding of the cerebral cortex (Zilles et al., 2013). For example, while species that are commonly used in neuroscientific studies (e.g., mice, rats, and marmosets) have relatively smooth (lissencephalic) cerebral cortices, 60-70% of the human cerebral cortex is buried in sulci (**Figure 1**; (Zilles et al., 1988; Van Essen, 2007; Willbrand et al., 2023c; Ramos Benitez et al., 2024). There are also many other mammal species that have highly gyrified cortex such as elephants, cetartiodactyls, carnivores, and others (Kazu et al., 2014; Lyras et al., 2016). Further, an array of recent studies has identified sulcal features that appear to occur exclusively in the cerebral cortex of hominid primates (i.e., humans and great apes—chimpanzees, bonobos, gorillas, and orangutans) from (i) smaller and shallower cortical indentations that are present in association cortices that are often functionally, behaviorally, and translationally relevant to (ii) deep sulcal points (Yücel et al., 2002; Bogart et al., 2012; Borst et al., 2014; Garrison et al., 2015; Leroy et al., 2015; Amiez et al., 2018; Lopez-Persem et al., 2019; Weiner, 2019; Nakamura et al., 2020; Rollins et al., 2020; Ammons et al., 2021; Hopkins et al., 2021a; Miller et al., 2021; Voorhies et al., 2021; Harper et al., 2022; Willbrand et al., 2022b, 2024b; Yao et al., 2022; Parker et al., 2023; Maboudian et al., 2024). Here, we consider branches, or rami, in the superior temporal cortex (STC).

**Figure 1:**
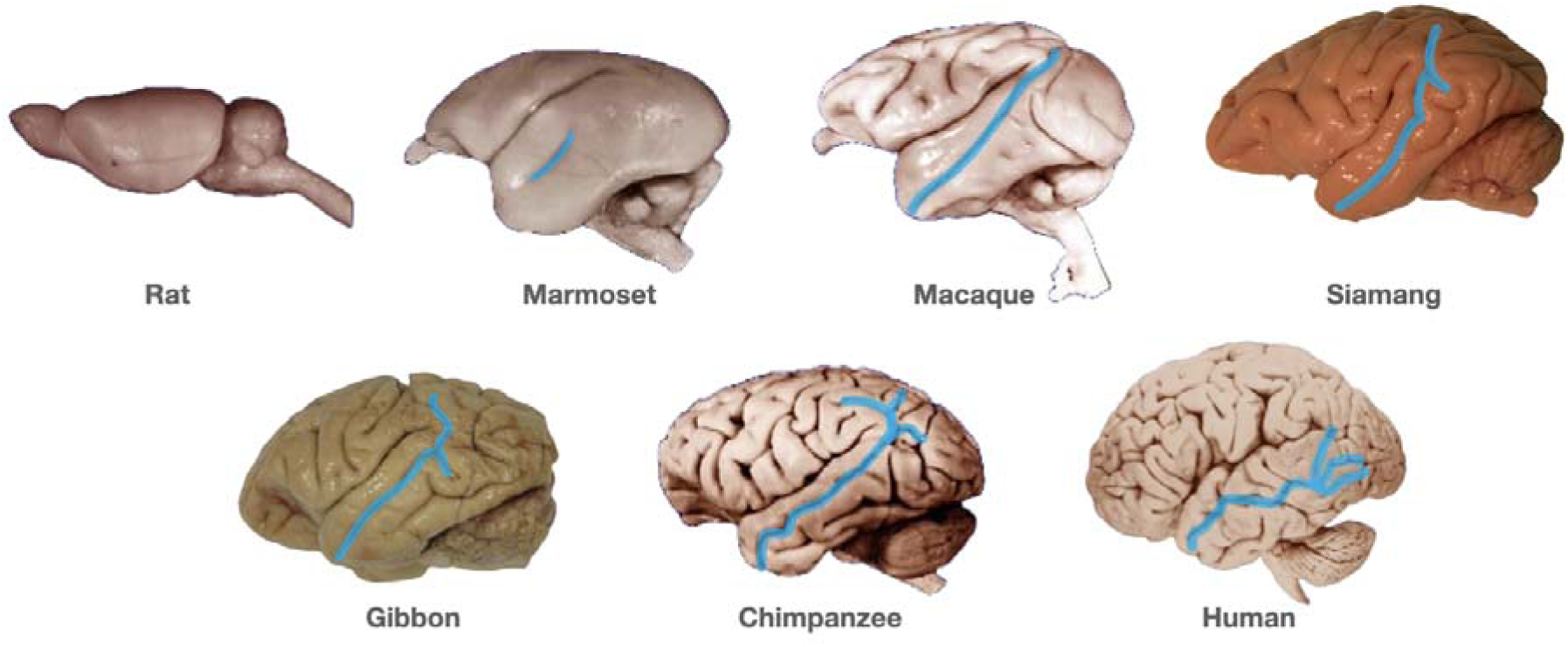
Example STS across mammalian brains. As pictured, rats do not have an STS, while marmosets, macaques, chimpanzees, and humans do have an STS. The macaque STS is prominent, although there are not three caudal rami. In siamang and gibbon, there are two caudal rami. In chimpanzees and humans, three caudal rami are present. Brain images are from https://brainmuseum.org/Specimens/primates/index.html and the archives of authors W.D.H. and C.C.S. Images not to scale.

Paleoanthropological and neuroimaging investigations show that STC is a cortical region that has expanded extensively throughout primate evolution (Segal and Petrides, 2012; Zilles et al., 2013; Leroy et al., 2015; Bruner, 2018; Van Essen et al., 2018; Petrides, 2019; Bruner et al., 2023; Willbrand et al., 2023d; Labra et al., 2024). Nevertheless, while the branches, or rami, of the superior temporal sulcus (STS), are specific to hominid brains (**Figure 1**) and have been consistently identified since the late 1800s across species (Kükenthal, 1895; Bolk, 1910; Shellshear, 1927; Connolly, 1950); **Figure 2**), the morphology of these branches has not been quantitatively compared between humans and great apes. Most recently, the STS has been referred to as having a “chaotic morphology,” especially in the left hemisphere (Le Guen et al., 2018), offering the intriguing opportunity to address this chaos between humans and chimpanzees.

**Figure 2.**
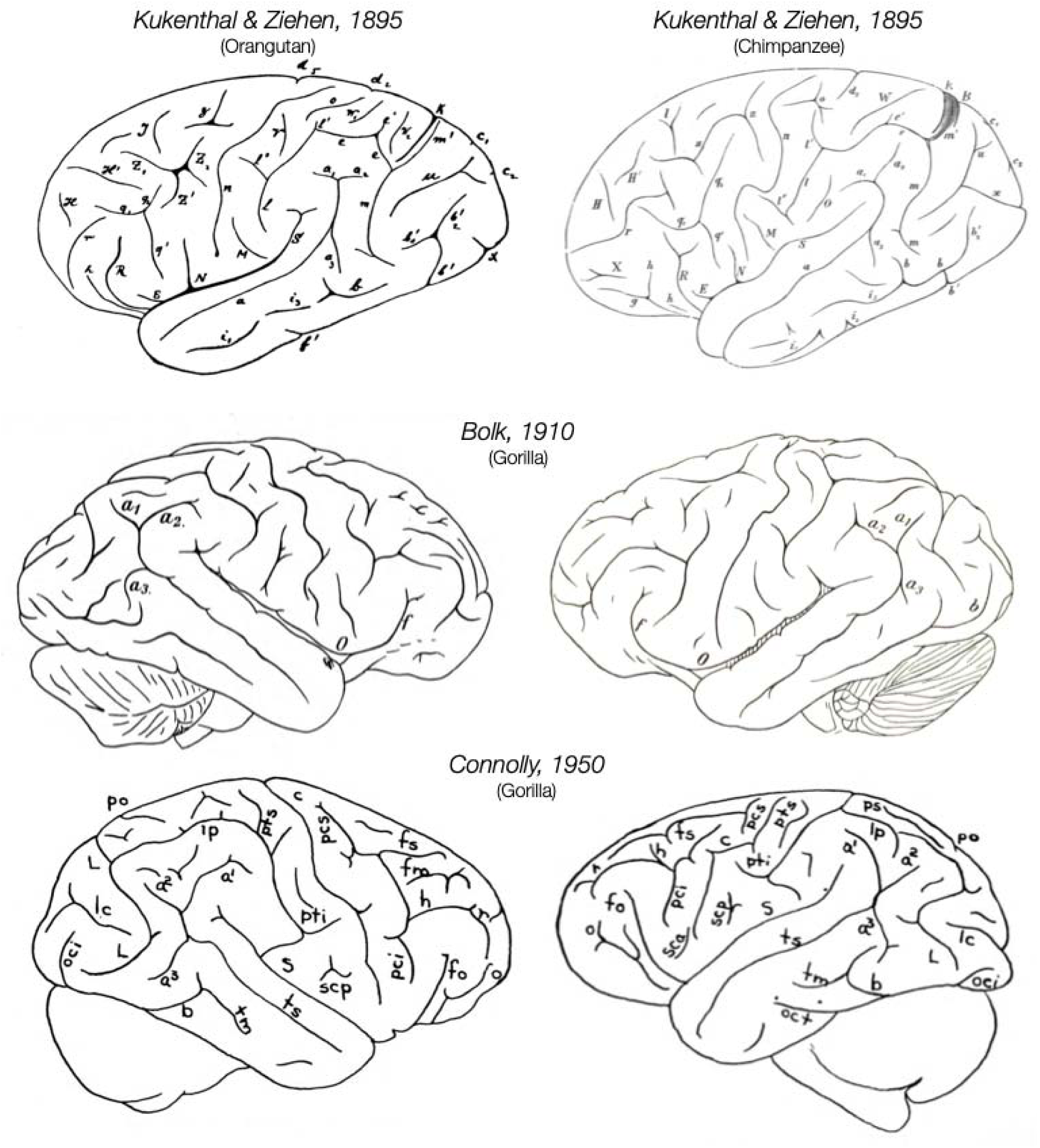
Three caudal rami of the STS in an orangutan, gorilla, and chimpanzee as depicted in historical sources. Though many modern neuroanatomical atlases and neuroimaging software packages identify a single STS in humans, neuroanatomists have labeled three caudal rami in great apes since the late 1800s. Historical labels of these rami (a1, a2, a3), anatomist, year, and species are included above in the images. Retzius (Retzius, 1906) should also be considered as he referenced posterior rami of the STS (specifically termed the “Ramus posterior sulci tempor. sup.”) which he identified as *ts’* (e.g., see Tables XLIV, XLVII, XLVIII, L, and LI).

To quantify this anatomy in STC between species, in the present study, we first sought to verify the occurrence of STS caudal rami in great ape brains and to comprehensively compare the morphology of the STS caudal rami between humans and chimpanzees, one of our species’ closest extant phylogenetic relatives (Schrago and Voloch, 2013). Given that features of the STS are cognitively and functionally relevant (Hein and Knight, 2008; Deen et al., 2015; Basil et al., 2017; Schobert et al., 2018; Bukowski and Lamm, 2020; Rollins et al., 2020; Lerosier et al., 2024), comparing the caudal rami of the STS between chimpanzees and humans will not only illuminate the comparative neuroanatomy of the STS between species, but also provide a foundation to further understand the evolutionary emergence of structural-functional and structural-behavioral relationships in this complex cortical expanse (Hein and Knight, 2008; Redcay, 2008; Hecht et al., 2013; Deen et al., 2015; Bruner, 2018; Specht and Wigglesworth, 2018; Bukowski and Lamm, 2020; Miller and Weiner, 2022; Bruner et al., 2023). Altogether, the present study aims to bridge the gap between historic qualitative observations and modern quantitative measurements in a part of the brain that has expanded substantially throughout evolution and that is involved in human-specific aspects of cognition.

## Materials and Methods

### Participants

#### Humans

Data from the young adult human cohort analyzed in the present study were from the Human Connectome Project (HCP) database (db.humanconnectome.org). Here, we used 72 participants (50% female; aged 22-36) that were also analyzed in several prior studies (Miller et al., 2020, 2021; Willbrand et al., 2022a, 2023a; b; c, 2024a; Hathaway et al., 2023; Maboudian et al., 2024), including a prior investigation of STC (Willbrand et al., 2023d).

#### Chimpanzees

Data from the adult chimpanzee cohort analyzed in the present study were from the National Chimpanzee Brain Resource (www.chimpanzeebrain.org). Here, we used 60 chimpanzees (63% female, aged 9-54) that were also analyzed in several prior studies (Miller et al., 2020; Willbrand et al., 2022a, 2023c; Hathaway et al., 2023). 30 chimpanzees were used to create a species-specific average template and were not included in any other analyses. Of the remaining chimpanzees, 29 were included in the manual labeling and analyses as one was excluded due to substantial issues in the cortical reconstruction pipeline. Chimpanzee MRIs were obtained from a data archive of scans collected prior to the 2015 implementation of U.S. Fish and Wildlife Service and National Institutes of Health regulations governing research with chimpanzees.

#### Bonobos, gorillas, and orangutans

Data from a cohort of nine adult or subadult great apes— three bonobos (*Pan paniscus*; two female), two gorillas (*Gorilla gorilla gorilla*; one female), and four orangutans (*Pongo pygmaeus*; one female)—were leveraged from prior work (Hopkins et al., 1998).

### Data Acquisition

#### Humans

Anatomical T1-weighted MRI scans (0.8 mm voxel resolution) were obtained in native space from the HCP database, along with outputs from the HCP modified FreeSurfer pipeline (Glasser et al., 2013).

#### Chimpanzees

Detailed descriptions of the scanning parameters have been described in Keller et al. (Keller et al., 2009), but we also describe the methods briefly here. Specifically, T1-weighted magnetization-prepared rapid-acquisition gradient echo (MPRAGE) MR images were obtained using a Siemens 3 T Trio MR system (TR = 2300 ms, TE = 4.4 ms, TI = 1100 ms, flip angle = 8, FOV = 200 mm) at the Emory National Primate Research Center (ENPRC) in Atlanta, Georgia. Before reconstructing the cortical surface, the T1 of each chimpanzee was scaled to the size of the human brain. As described in Hopkins et al. (Hopkins et al., 2017), within FSL, the BET function was used to automatically strip away the skull, (2) the FAST function was used to correct for intensity variations due to magnetic susceptibility artifacts and radio frequency field inhomogeneities (i.e., bias field correction), and (3) the FLIRT function was used to normalize the isolated brain to the MNI152 template brain using a 7 degree of freedom transformation (i.e., three translations, three rotations, and one uniform scaling), which preserved the shape of individual brains. Next, each T1 was segmented using FreeSurfer. The fact that the brains are already isolated, both bias-field correction and size-normalization, greatly assisted in segmenting the chimpanzee brain in FreeSurfer. Furthermore, the initial use of FSL also has the specific benefit, as mentioned above, of enabling the individual brains to be spatially normalized with preserved brain shape, and the values of this transformation matrix and the scaling factor were saved for later use.

#### Bonobos, gorillas, and orangutans

Again, detailed descriptions of the scanning parameters have been described elsewhere (Hopkins et al., 1998), but we also describe the methods briefly here. All procedures were conducted in accordance with protocols approved by the ENPRC. Subjects were initially immobilized via intramuscular injection of ketamine hydrochloride (10 mg/kg), followed by maintenance anesthesia using a continuous infusion of propofol (2–6 mg/kg/h). Animals were transported under anesthesia by van to the MRI facility at Emory University Hospital and remained anesthetized throughout transport and imaging procedures, which lasted approximately two hours in total. Post-scan, subjects were temporarily housed in single cages for 6–12 hours to allow recovery from anesthesia before being returned to their home enclosures and social groups. MRI scans were performed using two 1.5-Tesla Philips Model 51 scanners equipped with superconducting magnets. T1-weighted structural images were acquired in the transverse plane using a gradient echo protocol with the following parameters: repetition time = 19.0 ms, echo time = 8.5 ms, slice thickness = 1.2 mm, slice overlap = 0.6 mm, number of signal averages = 8, and matrix size = 256 × 256. These parameters were optimized based on preliminary studies and yielded high-resolution images suitable for morphometric analysis. Digital image data were archived onto optical diskettes and securely transferred for subsequent processing and analysis.

### Manually defining caudal rami of the STS in STC

#### Humans

We first manually defined the STC sulci within each individual hemisphere using *tksurfer* tools in FreeSurfer as described in our prior work (Miller et al., 2021). Manual lines were drawn on the inflated cortical surface based on the most recent schematics and studies of STC sulcal patterning (Segal and Petrides, 2012; Petrides, 2019; Willbrand et al., 2023d), as well as by the pial and *smoothwm* surfaces of each individual. Using the inflated, pial, and *smoothwm* surfaces to inform our labeling allowed us to form a consensus across surfaces and clearly determine each sulcal boundary.

Sulci were manually defined using guidance from the most recent atlas by Petrides (Petrides, 2019), as well as recent empirical studies (Segal and Petrides, 2012; Willbrand et al., 2023d), which together offer a comprehensive definition of cerebral sulcal patterns, including evolutionarily new sulci. The cortical region of interest was bounded by the following sulci and gyri: (i) the postcentral sulcus (PoCS) served as the anterior boundary, (ii) the superior temporal sulcus (STS) served as the inferior boundary, (iii) the intraparietal sulcus (IPS) served as the superior boundary, and (iv) the transverse occipital sulci (TOS) served as the posterior boundary. In the present study, we specifically examined the three different branches of the caudal superior temporal sulcus (posterior to anterior: cSTS3, 2, 1; (Segal and Petrides, 2012; Petrides, 2019). For each hemisphere, the location of the three cSTS rami were confirmed by trained independent raters (E.H.W., Y.T., T.G.) and finalized by a neuroanatomist (K.S.W.). The morphological features of these sulci in humans have already been published (Willbrand et al., 2023d), but have yet to be compared across species, which was the goal of the present study. See **Figure 3A** (right) for an example human hemisphere with the cSTS defined. All human cSTS sulcal definitions are in Supplemental Materials of our prior work (Willbrand et al., 2023d).

**Figure 3.**
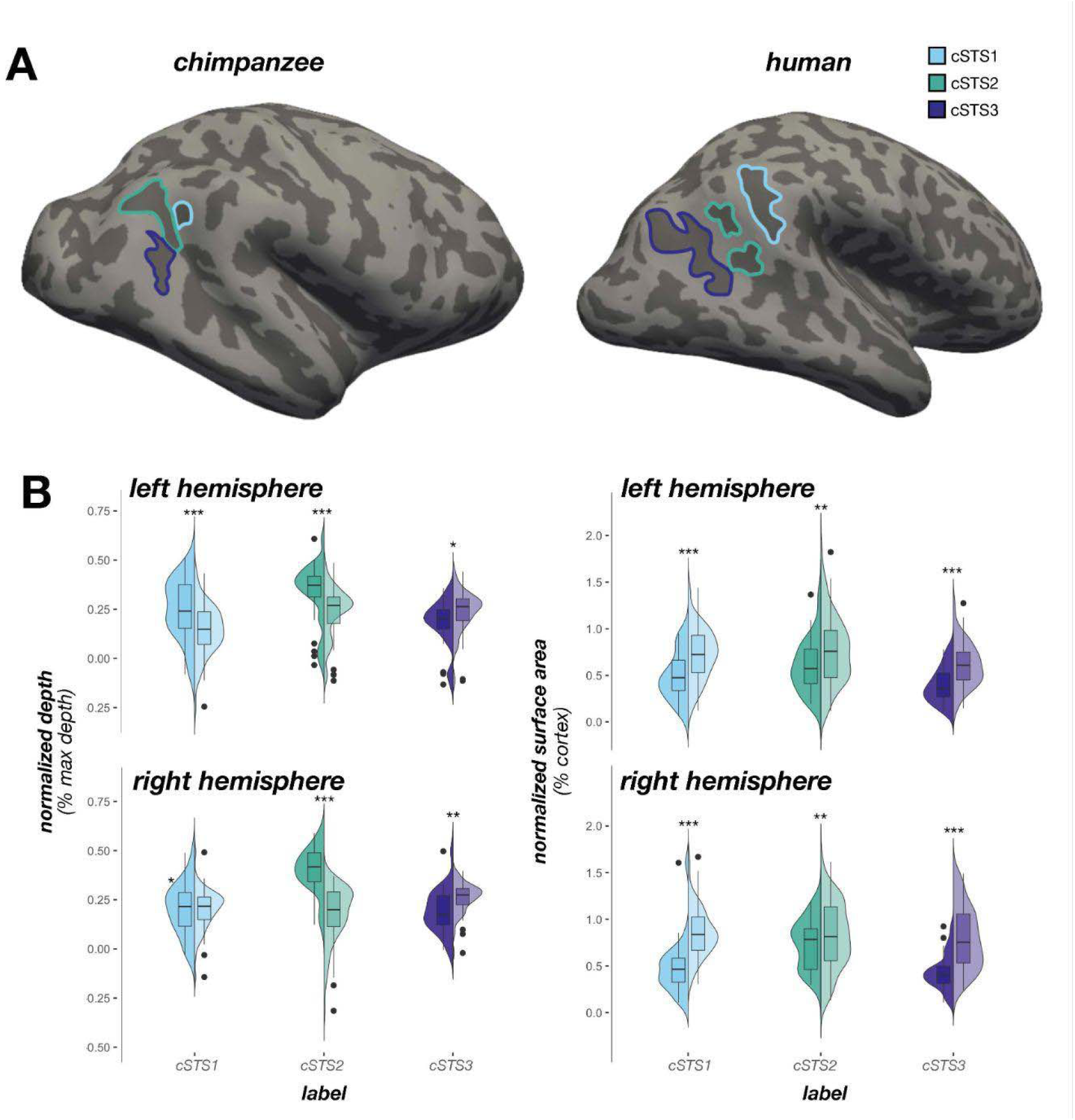
The caudal rami of the STS in chimpanzees. **A.** Chimpanzee (left) and human (right) inflated cortical surface reconstructions with the three caudal rami of the STS (cSTS) outlined on each surface (key). Sulci: dark gray; Gyri: light gray. **B.** Left: Split violin plots (box plot and kernel density estimate) visualizing normalized sulcal depth (percent of max depth; percentage values are out of 100) as a function of sulcus (x-axis), species (darker colors, right violin: human; lighter colors, left violin: chimpanzee), and hemisphere (top: left hemisphere; bottom: right hemisphere). Significant differences between species (as a result of the species x sulcus x hemisphere interaction) are indicated with asterisks (*p < .05, **p < .01, ***p < .001). Right: Same as left, but for normalized surface area (percent of parietal surface area; percentage values are out of 100). Significant differences between species (as a result of the species x sulcus interaction) are indicated with asterisks (**p < .01, ***p < .001).

### Great apes

The same sulci were also identified in chimpanzees, bonobos, gorillas, and orangutans when present. As with humans, for each hemisphere, the location of STC sulci was confirmed by trained independent raters (R.N.S., W.I.V., P.N.) and finalized by a neuroanatomist (K.S.W.). See **Figure 3A** (left) for an example chimpanzee hemisphere with the cSTS defined. All chimpanzee cSTS sulcal definitions are in **Supplemental Figure 1**. Given the smaller sample sizes of the bonobos, gorillas, and orangutans, we only quantified the incidence of the cSTS and did not include them in subsequent statistical analyses.

### Analyzing differences in sulcal incidence

All statistical analyses were implemented in R. We characterized the frequency of occurrence of each sulcus separately for left and right hemispheres. In line with prior work (Amiez et al., 2019, 2021; Willbrand et al., 2023c), for any sulcus that was variably present in either species, we tested the influence of species and hemisphere on the probability of a sulcus to be present with binomial logistic regression GLMs. For each statistical model, species (human, chimpanzee) and hemisphere (left, right), as well as their interaction, were included as factors for presence (absent, present) of a sulcus. Analysis of variance (ANOVA) chi-squared (*χ*2) tests were applied to each GLM, from which results were reported. GLMs were carried out with the glm function from the stats R package and ANOVA *χ*2 tests were carried out with the Anova function from the car R package.

### Extracting sulcal morphology

#### Depth

As in our prior work (Voorhies et al., 2021), mean sulcal depth values (in standard FreeSurfer units) were computed in native space from the .*sulc* file generated in FreeSurfer (Dale et al., 1999) with custom Python code. Briefly, depth values are calculated based on how far removed a vertex is from what is referred to as a “mid-surface,” which is determined computationally so that the mean of the displacements around this “mid-surface” is zero. Thus, generally, gyri have negative values, while sulci have positive values. To address scaling concerns between species, as in prior work (Miller et al., 2020; Hathaway et al., 2023; Willbrand et al., 2023c), each depth value was normalized by the deepest point in the given hemisphere (i.e., the insula).

#### Surface area

Surface area (mm^2^) was generated for each sulcus from the *mris_anatomical_stats* function in FreeSurfer (Dale et al., 1999; Fischl et al., 1999). To address scaling concerns between species, as in prior work (Miller et al., 2020; Hathaway et al., 2023; Willbrand et al., 2023c), we report surface area relative to the total surface area of the respective lobe that these sulci reside in (parietal lobe).

### Morphological comparisons

All morphological comparisons were conducted using linear mixed effects models (LMEs). ANOVA F-tests were subsequently applied. For both depth and surface area analyses, model predictors included sulcus, hemisphere, and species, as well as their interaction terms. Species, hemisphere, and sulcus were considered fixed effects. Sulcus was nested within hemisphere which was further nested within subjects. LMEs were implemented with the lme function from the nlme package. ANOVA F-tests were run with the aov function from the stats R package. Post-hoc analyses were computed with the emmeans and contrast functions from the emmeans R package. Post hoc p-values were corrected with the Tukey multiplicity adjustment. In the present study, we focus on species-related effects given that the morphological comparisons between these sulci were already documented by our group in the human sample (Willbrand et al., 2023d).

### Probability maps

Sulcal probability maps were calculated to summarize those vertices that had the highest and lowest correspondence across individual chimpanzees, respectively. To generate these maps, each sulcal label was transformed from the individual to a chimpanzee template surface from a held-out population of 30 chimpanzee brains that was made with the FreeSurfer *make_average_subject* function (Miller et al., 2020). Once transformed to this common template space, for each vertex, we calculated the proportion of chimpanzees for whom the vertex is labeled as the given sulcus. In the case of multiple labels, we employed a greedy, “winner-take-all” approach such that the sulcus with the highest overlap across participants was assigned to a given vertex. In addition to providing unthresholded maps, we also constrain these maps to maximum probability maps (MPMs) with 20% participant overlap as MPMs help to avoid overlapping sulci and increase interpretability (Miller et al., 2021). Human STC sulcal probability maps are available through our prior work (Willbrand et al., 2023d).

## Results

We first sought to identify the three caudal rami of the STS (cSTS) in chimpanzees, and then compare the incidence and morphology of these sulci to humans. Segal and Petrides (Segal and Petrides, 2012) have an excellent section in their paper describing the variability of how these three branches have been referenced throughout the classic literature from Smith (Smith, 1907), Economo and Koskinas (von Economo and Koskinas, 1925), Shellshear (Shellshear, 1927), Cunningham (Cunningham, 1931), Ono and colleagues (Ono et al., 1990), and Duvernoy (1999; please refer to Table 1 in Segal and Petrides (Segal and Petrides, 2012) and refer to the Supplemental Materials here for direct quotations from these authors). Nevertheless, those studies are largely restricted to the human cerebral cortex (though Shellshear (Shellshear, 1927) focuses on comparative analyses across species), and due to the high incidence rates of these caudal rami in chimpanzees, we returned to classic neuroanatomical texts to explore if these three rami were identifiable in great apes by prior neuroanatomists. Several different neuroanatomists identified and labeled these three cSTS branches in orangutans, gorillas, and chimpanzees (**Figure 2**). This historical classification was replicated in our chimpanzee sample as the three cSTS rami were identifiable in all chimpanzee cortical surface reconstructions (N = 29, 58 hemispheres; exemplary hemisphere in **Figure 3A**; all hemispheres in **Supplemental Figure 1**). We also found that the three cSTS rami were present in cortical surface reconstructions of other hominid species (bonobos, N = 3, 6 hemispheres; gorillas, N = 2, 4 hemispheres, orangutans, N = 4, 8 hemispheres; see **Figure 4** for example hemispheres) from previously published work (Hopkins et al., 1998). Finally, we found that the three cSTS rami were variably present in gibbon and siamang postmortem brains (**Supplemental Materials**).

**Figure 4.**
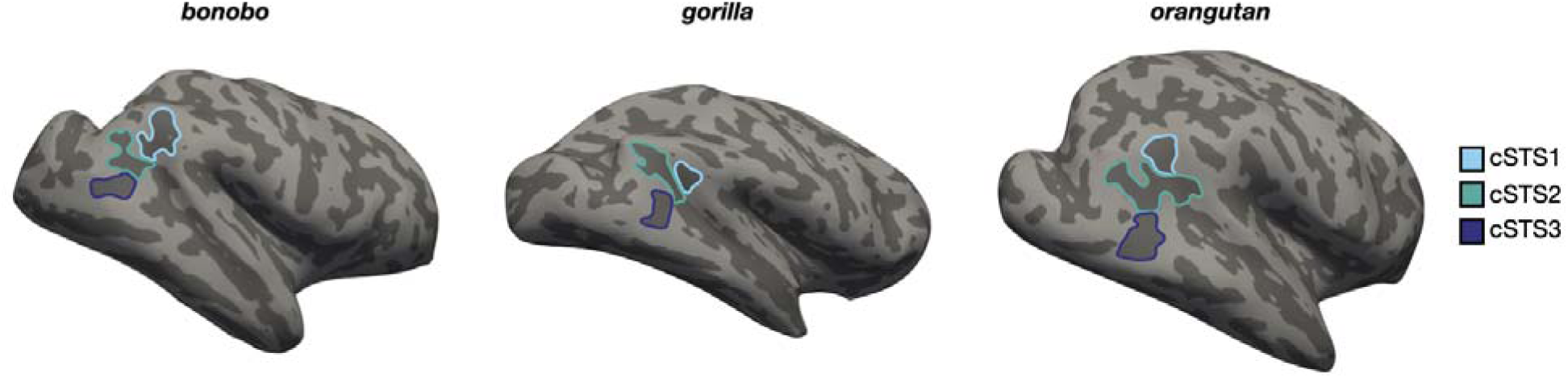
The caudal rami of the STS are present in three additional hominid species. Bonobo (left), gorilla (middle), and orangutan (right) inflated cortical surface reconstructions with the three caudal rami of the STS (cSTS) outlined on each surface (key). Sulci: dark gray; Gyri: light gray.

We next sought to examine if the three cSTS rami differed morphologically between humans and chimpanzees. In terms of depth (normalized to maximum hemispheric depth), an LME with predictors of sulcus, hemisphere, and species revealed three species-related findings. First there was a main effect of species (F(1, 593) = 23.26, p = 1.8 x 10^-6^), such that the cSTS components were relatively deeper in chimpanzees (**Figure 3B**, left). Second, there was a species x sulcus interaction (F(2, 593) = 36.65, p = 9.87 x 10^-16^) which showed that cSTS1 and cSTS2 were relatively deeper in chimpanzees (ps < .004), whereas cSTS3 was relatively deeper humans (p = .001; **Figure 3B**, left). Finally, there was a species x sulcus x hemisphere interaction (F(2, 593) = 9.98, p = 5.45 x 10^-5^). Post hoc pairwise comparisons revealed that: (i) cSTS1 was deeper in chimpanzees in the left (p < .0001), but not right (p = .94) hemisphere, (ii) cSTS2 was deeper in chimpanzees in both hemispheres (ps < .001), and (iii) cSTS3 was deeper in humans in both hemispheres (ps < .05; **Figure 3B**, left). In terms of surface area (normalized to hemispheric parietal lobe surface area), a similarly-structured LME revealed two species-related findings. First, there was a main effect of species (F(1, 593) = 83.97, p < 2 x 10^-16^) such that the cSTS components were relatively larger in humans (**Figure 3B**, right). Second, there was a species x sulcus interaction (F(2, 593) = 4.81, p = .008) where post hoc pairwise comparisons identified that cSTS1 and cSTS3 exhibited larger differences in size between species (ps < .0001) than cSTS2 (p = .006; **Figure 3B**, right). There was no species x sulcus x hemisphere interaction (p = .12; **Figure 3B**, right). To ensure that our results were not simply due to normalization, we also implemented the same analysis on the raw quantitative morphological metrics which confirmed these findings (**Supplemental Materials**).

## Discussion

By manually defining STC sulci in 144 human and 58 chimpanzee hemispheres, we show that the surface anatomy of this cortical expanse is both similar and different between these two closely related hominid species along three sulcal metrics: i) incidence, ii) depth, and iii) surface area. These findings demonstrate several key similarities and differences that shed light on the evolutionary trajectory of cortical folding in this region.

First, we found that all three cSTS branches were identifiable in every chimpanzee hemisphere as well as the other great apes brains we examined, mirroring the consistent presence previously documented in human populations (Segal and Petrides, 2012; Willbrand et al., 2023d). This high incidence rate across species *suggests* that these sulci are likely homologous and evolutionarily conserved features of hominid brains. Supporting this conclusion, we found similar anatomical positioning of these sulci in both humans and chimpanzees, consistent with classic schematics proposed by neuroanatomists over a century ago (e.g., (Shellshear, 1927; Connolly, 1950); Figure 1, **Supplementary Materials**). These results contribute to a growing body of evidence (e.g., (Amiez et al., 2019, 2021, 2023; Miller et al., 2020; Hopkins et al., 2022; Willbrand et al., 2022a, 2023c; Hathaway et al., 2023) showing that while some sulci may be evolutionarily novel in humans, others—even in association cortices—have evolutionary roots shared with great apes, with potential functional insights. For instance, previous work shows that particular branches of the human STS predict the location of functional regions involved in face processing that have a consistent topological organization relative to functional regions selective for bodies and visual motion in a cortical expanse now referred to as the “lateral” visual stream (Weiner and Grill-Spector, 2013; Gomez et al., 2019; Finzi et al., 2021; Pitcher and Ungerleider, 2021; Weiner and Gomez, 2021).

Despite the consistent presence of these three caudal rami in both humans and chimpanzees, we did identify quantitative morphological differences. Specifically, the cSTS branches were relatively deeper in chimpanzees, but had a greater surface area in humans. These evolutionary differences may reflect evolutionary pressures toward increased functional specialization in this part of the cortex, as depth and surface area are theorized to be linked to increased cortical surface available for local circuitry and potential expansion of specific functional features (Van Essen, 2007; Miller and Weiner, 2022). Indeed, neuroimaging studies have identified this region to be associated with complex higher-level cognitive functions (e.g., theory of mind, audiovisual integration, speech processing, and face processing)—functions thought to be associated with cortical areas that have expanded in humans relative to non-human primates (Hein and Knight, 2008; Redcay, 2008; Deen et al., 2015; Specht and Wigglesworth, 2018; Bukowski and Lamm, 2020). These differences may also reflect “compensatory” mechanisms. Specifically, Connolly’s compensation theory of cortical folding proposes that the quantitative features of sulci are likely counterbalanced by those of their neighbors (Connolly, 1940, 1950). In terms of the compensation theory, then, the decreased cSTS depth in humans may reflect the presence of an increased number of putative tertiary sulci apparently present in this cortical expanse in humans relative to chimpanzees, which can be tested in future studies.

Beyond macroanatomical comparisons, genetic and neurodevelopmental frameworks offer a critical context for interpreting species-specific sulcal morphology. For example, studies in both humans and chimpanzees have linked variation in STC morphology and function to polymorphisms in the *KIAA0319* gene, which is implicated in cortical development and associated with reading and language disorders in humans (Harold et al., 2006; Darki et al., 2012; Pinel et al., 2012; Jamadar et al., 2013; Hopkins et al., 2021b, 2023; Paniagua et al., 2022). Of direct relevance to the present study, one of these studies identified that the depth and surface area of the STS (especially in the central portion) were highly heritable and that polymorphisms in two genes (*KIAA0319* and *AVPR1A*) were associated with average STS depth and asymmetry (Hopkins et al., 2023). The presence of similar associations in chimpanzees and humans suggest a potentially conserved genetic influence on the organization of this region. Further, as the caudal rami of the STS examined in the present study are cortically distant from the most heritable portion of the STS identified by Hopkins et al. (Hopkins et al., 2023); i.e., the central portion), this distance may partially explain the interspecies differences in the cSTS identified in the present study, given that strongly heritable sulci have been shown to be more evolutionarily conserved in humans and chimpanzees (Gómez-Robles et al., 2015; Amiez et al., 2018; Pizzagalli et al., 2020; Schmitt et al., 2021; Hopkins et al., 2023). Human studies have also shown that the left posterior STS is particularly susceptible to morphological variability, with features such as “pli-de-passage” (buried gyri) and sulcal fragmentation occurring more frequently in the left hemisphere and under moderate genetic control (Le Guen et al., 2018; Bodin et al., 2021). Such variability may then reflect both genetic constraints and experience-dependent changes in cortical development.

Together, these lines of evidence underscore that morphological evolution in the STC likely results from an interplay of conserved genetic architecture and species-specific neurodevelopmental trajectories, which may link variations in sulcal anatomy to the emergence of complex functional and cognitive features. Future studies should directly assess the role of genetic variations (such as polymorphisms in the *KIAA0319* gene) as well as that of other non-genetic factors (such as age) in the stability of and changes in the morphology of all STC sulci across hominid species, and whether these features have differing impacts on the morphology of the STS and the three cSTS rami specifically.

Furthermore, our results suggest that while the qualitative sulcal presence of the cSTS branches in the STC is conserved across humans and great apes, alterations have occurred at the quantitative level. These changes, which are especially striking compared to other species (**Figure 1**), parallel the cognitive and behavioral evolution of humans, particularly in visuospatial, attentional, and tool-use domains. These findings lay the groundwork for future investigations that link sulcal morphology in the STC to individual differences in cognition and behavior—both within and across species—and offer new insights into the neural basis of human-specific cognitive capacities.

## Limitations and Future Directions

There are several limitations to this work. First, while our chimpanzee sample (N<50) is comparable to many nonhuman primate neuroimaging studies, it is still modest compared to large N (N>1,000) human neuroimaging datasets. Second, while we were able to compare sulcal incidence and morphology, we did not directly assess other relevant features (e.g., microanatomical, functional, or behavioral data) between species. Future analyses from rare functional MRI and behavioral data in chimpanzees could help clarify the extent to which sulcal landmarks correspond to functional representations within STC and behavior (as has been done in anterior cingulate cortex; (Amiez et al., 2021; Hopkins et al., 2021a). Additionally, developmental studies in both species could help determine whether differences in sulcal morphology arise from differences in developmental timing or rates of cortical expansion (Sakai et al., 2012). To aid the identification of these sulci in such future studies, we include chimpanzee probabilistic maps of these sulci (**Data accessibility statement**; **Figure 5**).

**Figure 5.**
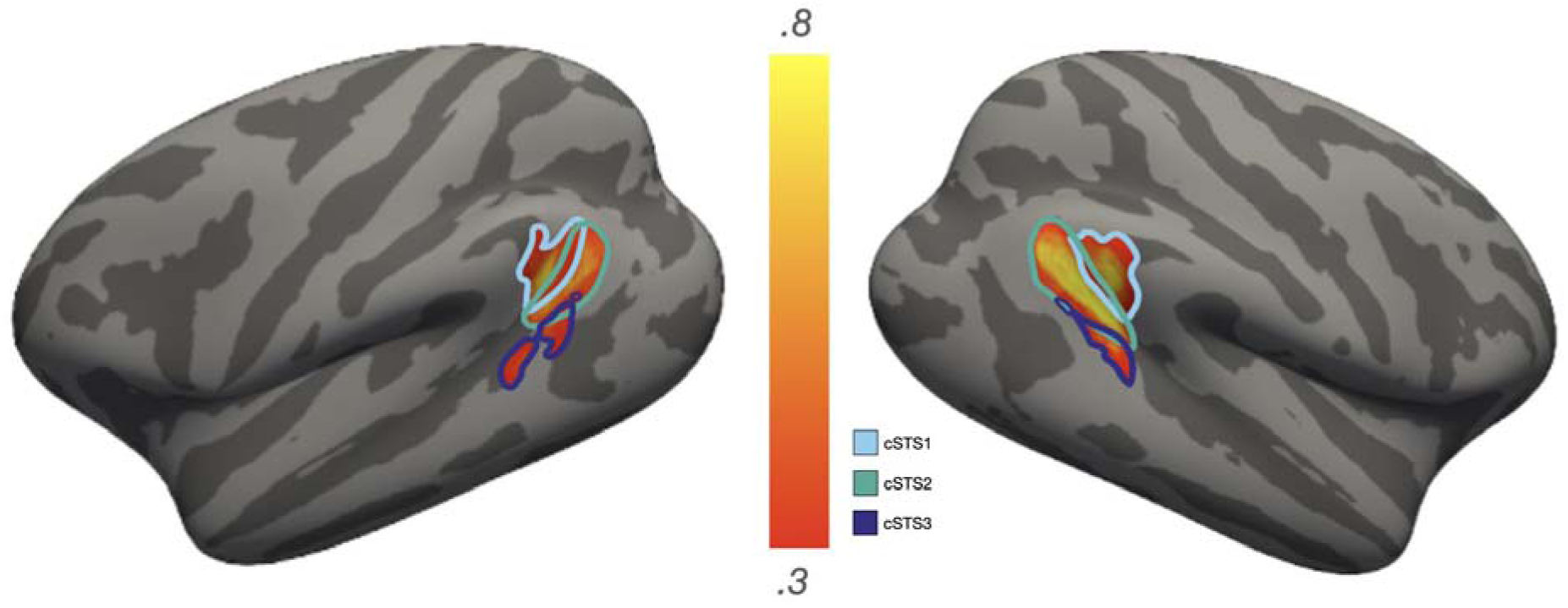
cSTS sulcal probability maps in chimpanzees. Probability maps for the three cSTS rami identified in the present work. To generate the maps, each label was transformed from each individual to a custom average template created from 30 additional chimpanzees that were not included in the original analysis. For each vertex, we calculated the proportion of chimpanzees for whom that vertex is labeled as the given sulcus (the warmer the color, the higher the overlap in each image). In the case of multiple labels for one vertex, the sulcus with the highest overlap across participants was assigned to a given vertex. To reduce spatial overlap for visualization purposes, these maps were thresholded to variable degrees of overlap across chimpanzees (scales).

## Conclusion

This study provides the first (to our knowledge) direct quantitative comparative analysis of the three caudal rami of the STS between humans and chimpanzees. Our results reveal a shared sulcal framework with evidence of morphological differences in the human brain from that of chimpanzees. These findings illuminate both conserved and species-specific features of cortical organization that may advance our understanding of the structural foundations of human cognitive evolution.

### Ethics statement

#### Human participants

HCP consortium data were previously acquired using protocols approved by the Washington University Institutional Review Board. Informed consent was obtained from all participants

#### Chimpanzee participants

The chimpanzees were all members of the colony housed at the Emory National Primate Research Center (ENPRC). All methods were carried out in accordance with ENPRC and Emory University’s Institutional Animal Care and Use Committee (IACUC) guidelines. Institutional approval was obtained prior to the onset of data collection.

## Data accessibility statement

Data, code, analysis pipelines, and sulcal probability maps, will be made publicly available on GitHub upon the publication of this paper at: https://github.com/cnl-berkeley/stable_projects.

