## Supplementary material for "A morphological comparison of the caudal rami of the superior temporal sulcus in humans, chimpanzees, and other great apes": SM

[1]*Cognitive Science Program, University of California, Berkeley, Berkeley, CA, USA*
[2]*Medical Scientist Training Program, School of Medicine and Public Health, University of Wisconsin–Madison, Madison, WI USA*
[3]*Barnard College, Columbia University, New York City, NY, USA*
[4]*Calbright College, Sacramento, CA, USA*
[5]*Department of Psychology, University of North Carolina at Chapel Hill, Chapel Hill, NC, USA*
[6]*Medical Science Faculty, University of Bordeaux, Bordeaux, France*
[7]*Department of Anthropology and Center for the Advanced Study of Human Paleobiology, The George Washington University, Washington, DC, USA*
[8]*Department of Comparative Medicine, The University of Texas MD Anderson Cancer Center, Bastrop, TX, USA*
[9]*Department of Psychology, University of California, Berkeley, Berkeley, CA, USA*
[10]*Helen Wills Neuroscience Institute, University of California, Berkeley, Berkeley, CA, USA*
[11]*Department of Neuroscience, University of California, Berkeley, Berkeley, CA, USA*
**Co-first author*

**Keywords***:* Comparative biology, Cortical folding, MRI, Morphometry, Neuroanatomy, Superior temporal cortex, Superior temporal sulcus

**Supplemental Text**

### Historical references to caudal rami of the superior temporal sulcus

Below we include text from previous neuroanatomical investigations identifying caudal branches of the superior temporal sulcus (cSTS).

***From Segal and Petrides** (2012):* “For example, schematic illustrations of the brain’s lateral surface in both Duvernoy (1999) and Ono et al. (1990) represent two cSTS branches within the IPL. The schematic diagram in Duvernoy (1999, p. 7) labels two branches of the cSTS as the ascending and the horizontal posterior segments, while Ono et al. (1990, Ch. 2, p. 16) identify two cSTS branches as the angular sulcus and the anterior occipital sulcus. At first, it may seem that the same two branches are represented in these two atlases but under different names; closer examination indicates otherwise. The cSTS branch labeled as ascending by Duvernoy (1999) lies immediately behind the posterior ascending ramus of the Sylvian fissure (ascSF; see Duvernoy, 1999; sagittal sections, pp. 258–259). However, in Ono et al. (1990) a similar sulcus found just behind the ascSF is treated as an infrequent configuration of the STS and called the double parallel-type termination (Ch. 10, p. 77). The angular sulcus of Ono et al. (1990) now appears to refer to the horizontal posterior segment of Duvernoy. Thus, the above two atlases refer to two cSTS branches but they do not seem to identify the same two sulci (see Table 1 for summary).”

***From Shellshear** (1927):* “In Figure 9 the lower end of the sulcus occipitalis anterior is joined to a well developed sulcus temporalis medius. The usual description given in textbooks, originated without due regard to morphological facts, could be applied to this specimen. The sulcus temporalis superior is most commonly described as being the homologue of the parallel sulcus and ending in the angular gyrus. The middle temporal sulcus is diagrammatically represented as lying parallel with the sulcus temporalis superior and ending in the gyrus postparietalis. The detached portion of the superior parallel sulcus is disregarded or unrecognised. It is now clear that the superior temporal sulcus so described is a complex sulcus consisting of the anterior temporal (related to the area temporalis polaris), the inferior parallel (related to areas 21 and 22), a portion of the superior parallel and the angular sulcus (related to area 39), with in some cases-depending upon the point of junction of the sulcus angularisthe sulcus annectans. Further, when the angular sulcus is separate, the upper segment of the anterior occipital sulcus may be included. The middle temporal sulcus so described consists of a series of disconnected sulci (the proper middle temporal sulcus), occasionally joined together, connecting with the lower segment of the anterior occipital sulcus and thence continued to the postparietal lobule."

***From Bailey et al.** (1950):* "*ts, the sulcus temporalis superior*, *sulcus parallelis*. With Shellshear (1927) the parallel sulcus can be divided into an anterior and a posterior part. The former, one of the oldest furrows of the primate brain, runs in the temporal lobe; the latter, changing profoundly from macaque to man, runs in the parietal lobe. Near the temporal pole, the anterior end may be in line with the rest of the sulcus or may be bent ventrad so that the temporal pole appears as a continuation of the superior temporal gyrus. Occasionally, as Blinkow (1938) observed, this anterior hook may be an independent sulcus and the temporopolar region may be opercularized. Submerged bridging convolutions within the anterior part have been described by Blinkow. Within the angular gyrus the posterior part generally breaks up into three rami. Connections with the sulci of both occipital lobe and inferior parietal lobule are frequent. Shellshear reports that an interruption of the parallel sulcus between anterior and posterior part is not infrequent. No anthropological observations about this sulcus were found by us."

***From Tamraz and Comair** (2000):* "The posterior part is the angular sulcus which penetrates into the inferior parietal lobule and usually divides into three rami within the angular gyrus."

***From Connolly** (1950):* "THE ANTERIOR OCCIPITAL of Wernicke (a3) separates the gyrus angularis from an area caudal to it and termed by Brodmann (1925) area praeoccipitalis (area 19). As the sulcus is not within the occipital lobe, the term used by Genna (1924) namely s. preoccipitalis seems more appropriate than anterior occipital. Eberstaller (1884) described a sulcus which is axial to the postparietal part of the inferior parietal lobule and viewed it as an ascending ramus of the temporal medius sulcus which is apparently identical with the anterior occipital sulcus of Wernicke. Kohlbrugge (1909) regarded the sulcus as independent of the midtemporal and a doubling of the superior temporal sulcus. Shellshear (1927) gives a similar view. An upper and a lower part of the anterior occipital are, according to this view, both developed from the posterior wall of the superior parallel sulcus and at least in simply fissurated brains, are connected with it by the sulcus annectans. Wang and Kappers (1924) designate the anterior occipital or s. preoccipitalis as ascending ramus III, the superior parallel being the ascending ramus I and the s. angularis, the ascending ramus II. According to these authors the ascending branches are split off from ts."

**Raw cSTS depth and surface area compared between chimpanzees and humans**

In terms of depth (mm), a linear mixed effects model (LME) with predictors of sulcus, hemisphere, and species revealed three species-related findings. First there was a main effect of species ($F(1, 582) = 153.66$, $p = 2 \times 10^{-16}$), such that the cSTS components were deeper in chimpanzees (**Supplemental Figure 2A**). Second, there was a species x sulcus interaction ($F(2, 582) = 18.87$, $p = 1.14 \times 10^{-8}$) which showed that all branches were deeper in chimpanzees ($ps < .0007$; **Supplemental Figure 2A**). Finally, there was a species x sulcus x hemisphere interaction ($F(2, 582) = 6.542$, $p = .0015$) indicating similar relationships observed in the main text (**Supplemental Figure 2A**). In terms of surface area ($mm^2$), a similarly-structured LME revealed two species-related findings. First, there was a main effect of species (F(1, 582) =

121.86, $p < 2 \times 10^{-16}$) such that the cSTS components were larger in humans (**Supplemental Figure 2B**). Second, there was a species x sulcus interaction ($F(2, 582) = 3.95$, $p = .0019$) where post hoc pairwise comparisons showed that all three branches had a larger surface area in humans compared to chimpanzees ($ps < .0001$). There was no species x sulcus x hemisphere interaction ($p = .19$; **Supplemental Figure 2B**).

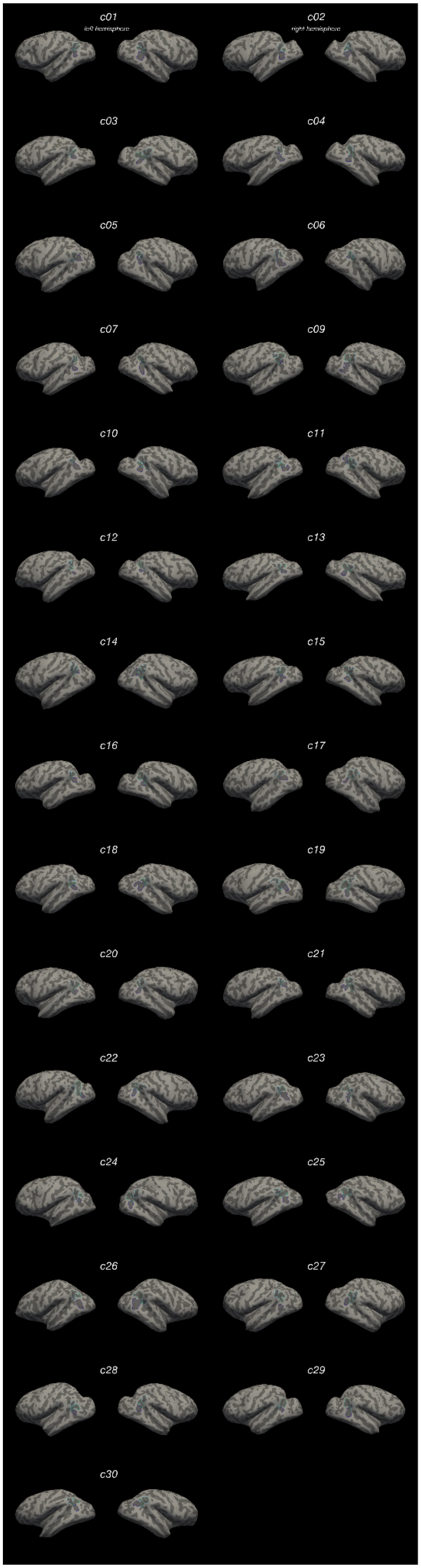
c01
left hemisphere
c02
right hemisphere
c03
c04
c05
c06
c07
c09
c10
c11
c12
c13
c14
c15
c16
c17
c18
c19
c20
c21
c22
c23
c24
c25
c26
c27
c28
c29
c30

**Supplemental Figure 1. Caudal rami of the superior temporal sulcus identified in every chimpanzee hemisphere.** Each sulcus is displayed on the left and right hemisphere inflated cortical surfaces in FreeSurfer 6.0.0, with the label displayed as an outline according to the key at the top.

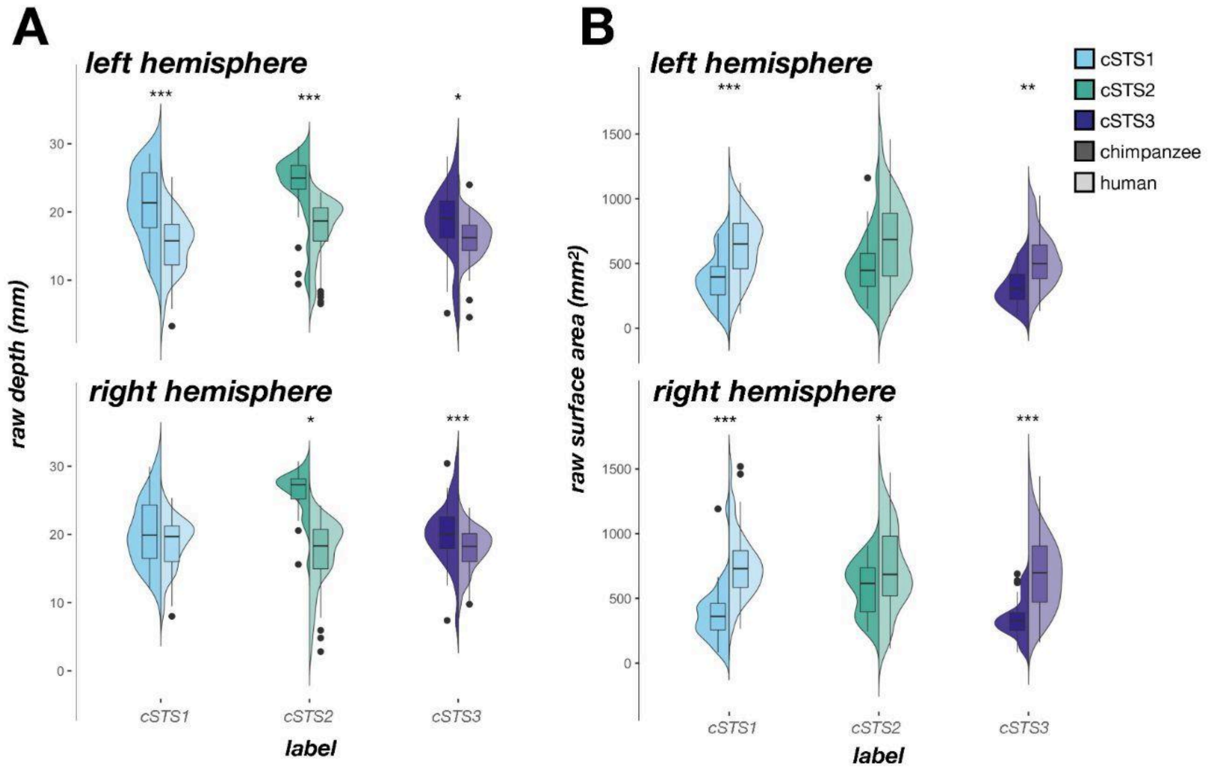


**Supplemental Figure 2. Raw depth and surface area of the caudal rami of the superior temporal sulcus in humans and chimpanzees. A.** Split violin plots (box plot and kernel density estimate) visualizing raw sulcal depth (mm) as a function of sulcus (x-axis), species (darker colors, right violin: human; lighter colors, left violin: chimpanzee), and hemisphere (top: left hemisphere; bottom: right hemisphere). Significant differences between species (as a result of the species x sulcus x hemisphere interaction) are indicated with asterisks (*$p < .05$, **$p < .01$, ***$p < .001$). **B.** Same as left, but for raw surface area ($mm^2$). Significant differences between species (as a result of the species x sulcus interaction) are indicated with asterisks (** $p < .01$, *** $p < .001$).

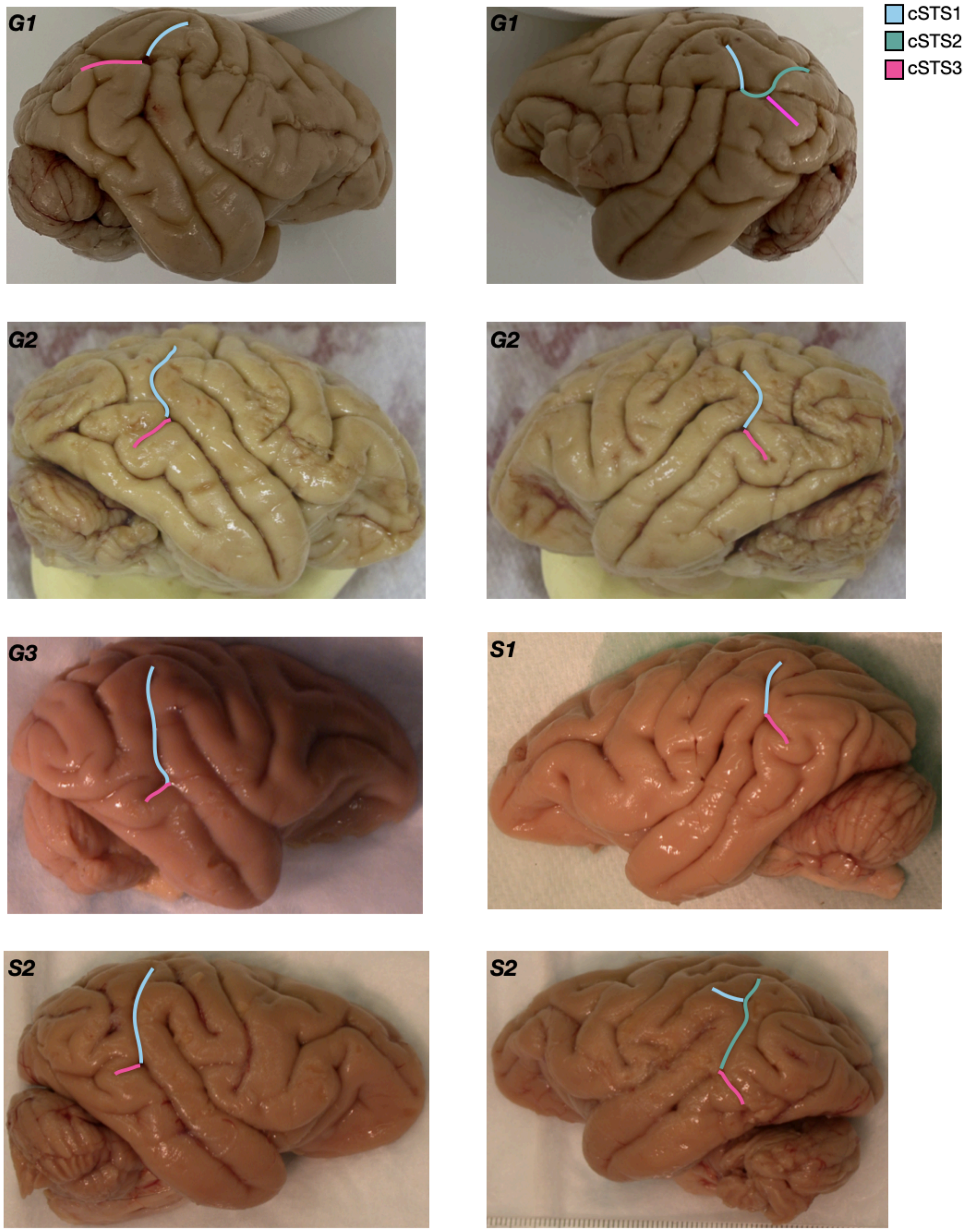


**Supplemental Figure 3. The caudal rami of the STS are variably present in gibbon and siamang postmortem brain images.** Gibbon (labeled G#) and siamang (labeled S#) postmortem brain images with the cSTS identified when present (key). Left hemispheres (LH) and right hemispheres (RH) are identified with their respective abbreviation. Brain images are from the archives of authors W.D.H. and C.C.S. Images not to scale.
